# Additive baselines furnish no evidence for epistasis learning by MULTI-evolve

**DOI:** 10.64898/2026.04.23.719915

**Authors:** Gian Marco Visani, Aayush Verma, William S. DeWitt

## Abstract

Recent work from Tran *et al*. (*Science*, 2026) introduced MULTI-evolve, a framework for protein engineering that combines single-mutant nomination via a protein language model (PLM) or a deep mutational scan (DMS), experimental single- and double-mutant characterization, and neural networks to engineer hyperactive multimutant proteins. The authors attribute the framework’s performance to “epistasis-aware modeling” and claim that their neural networks “learn the epistatic landscape” and “identify synergistic interactions” from limited double-mutant training data. Additive models, by definition, cannot represent epistasis, making them a natural null baseline for such claims. Here we show that MULTI-evolve’s multimutant predictions are almost perfectly correlated with an additive model’s across all three engineering applications (APEX, dCasRx, and HuABC2), such that the engineering of multimutants reduces to combining beneficial mutations with the largest additive effects—a standard protein engineering strategy for over four decades. We also find that MULTI-evolve’s neural networks do not outperform an additive model on held-out test set predictions, and do not even represent epistasis in their training data. Finally, we revisit a DMS benchmark finding presented as evidence of epistasis learning and show that the same pattern is expected even under a null additive model, due to an elementary statistical phenomenon; when we fit an additive model to the benchmark data, it reproduces the reported pattern. More broadly, our findings underscore the need to benchmark models for machine learning-guided directed evolution against additive null baselines before attributing performance to learned epistasis.

## Introduction

Tran *et al*. [1] present MULTI-evolve as a framework whose success hinges on learning epistatic interactions between mutations. Epistasis—the dependence of one mutation’s effect on the presence of others [2–4]—is invoked throughout the paper as both the challenge and the solution. The abstract describes “epistatic modelling to predict synergistic combinations.” The central computational claim is that “the predictive power of these models depends on learning epistatic effects in the target protein, where the functional impact of one amino acid mutation can alter others”. The authors “hypothesized that double mutants capture sufficient epistatic information for accurate extrapolation to higher order mutants”, and in the Discussion characterize their contribution as “a machine learning-guided framework that overcomes the challenge of a massive search space for protein engineering by bridging evolutionary PLM priors with epistatic modelling to access synergistic higher-order combinations”.

This language situates MULTI-evolve within a broader movement in machine learning-guided directed evolution (MLDE). Directed evolution has a rich experimental tradition [5], culminating in the 2018 Nobel Prize in Chemistry. More recently, ML-augmented approaches have sought to accelerate the process by training models on measured protein phenotypes and using them to propose untested variants [6–9]. A recurring claim in this literature is that ML models can learn epistatic interactions and thereby navigate rugged genotype-phenotype maps that defeat simpler strategies [10]. MULTI-evolve follows this pattern, asserting that its fully connected neural networks (FCNNs) “learn the epistatic landscape from a compact dataset of double mutants and extrapolate to synergistic combinations”.

The classical protein engineering literature provides a clear baseline against which to evaluate such claims. As reviewed by Wells in 1990 [11], it has long been recognized that mutational effects on protein function are often approximately additive—particularly on a logarithmic scale for independent thermodynamic contributions to stability or catalysis. Additivity is expected on biophysical grounds when mutations affect structurally independent contacts. The additive stacking strategy—identifying beneficial single mutations and combining them—is often productive because additivity often holds at least approximately [11–13]. A defining feature of an additive landscape is that interactions between mutations are identically zero, so benchmarking more complex nonlinear models against a standard additive model is critical for establishing that epistatic interactions and synergistic effects have been genuinely ascertained. Because no such comparison is presented in [1], we undertook it ourselves, finding no evidence to substantiate the claims that MULTI-evolve’s neural networks learn epistatic interactions. We propose a more parsimonious interpretation of MULTI-evolve’s performance, organized around a central finding: *MULTI-evolve’s neural network learns an additive model*. We hasten to add that this is a claim about MULTI-evolve, not about epistasis in general: epistasis is well-documented in protein genotype-phenotype maps [3, 14], and may in principle be learnable by ML models, but we will show that in the settings considered by [1] the FCNN collapses to an additive model with no learned epistatic interactions.

First, we consider the task of engineering higher-order mutants of APEX, dCasRx, and HuABC2 using the MULTI-evolve FCNN and comparing its predictions against an additive model across the combinatorial space. For each engineering task, we trained the FCNN and the additive model on the same training data (single and double mutants), and then generated predictions for all 5–9-mutant APEX variants, all 5–7-mutant dCasRx variants, and all 3–7-mutant HuABC2 variants (for both expression and binding phenotypes). We found that the FCNN’s predictions are almost perfectly correlated with the additive model’s predictions, such that the ranking of tens of thousands of untested variants is practically identical to an additive model. In particular, the multimutant variants selected for production are similarly highly ranked by both the FCNN and the additive model, calling into question the central claim that MULTI-evolve’s success is due to learning synergistic interactions and traversing a rugged epistatic landscape. For HuABC2, variants were selected along the expression–binding Pareto frontier; we show that the Pareto frontiers under the additive model and MULTI-evolve are nearly identical. We also find that the FCNN does not represent epistasis in its own training data. The engineering results are indistinguishable from additive stacking of highly beneficial single mutations, a strategy that has been standard in protein engineering for over four decades [11–13].

Second, we reran the MULTI-evolve hyperparameter grid search on the published APEX, dCasRx, and HuABC2 training data [15], but extended the grid to include zero hidden layers, tantamount to a linear model, and also compared to a classic ridge estimator of an additive model (both these extensions have no capacity to represent epistasis). We found that the nonlinear FCNNs of MULTI-evolve did not significantly improve on these non-epistatic baselines in terms of predictive performance on held-out test data.

Finally, we revisit a key deep mutational scan (DMS) [16] benchmarking result presented as evidence of MULTI-evolve’s epistasis learning capacity, and show that the same pattern is expected under a null additive model. The authors trained their FCNN on three nested subsets of sequences from previously published DMS data [14, 17–19]—single mutants only, single and double mutants only, and up to triple mutants—and observed improved prediction on held out multimutant data as the training set enlarged. We show that this pattern is expected even under a null model of additivity due to the elementary statistical effect of prediction error variance reduction with larger sample sizes. When mutational effects are additive, each double or triple mutant provides a redundant measurement of the same single-mutation coefficients, improving estimation without teaching the model anything about interactions. The paired *t* tests presented in [1, Fig. 2E] to quantify the signal of epistasis learning do not control for the variance reduction effect. To confirm this issue in the context of the DMS datasets (which likely do contain pervasive epistatic effects), we fit an additive model on the same nested DMS data MULTI-evolve used, and reproduced the pattern in [1, Fig. 2E]. Furthermore, we question the relevance of this benchmarking procedure, which trains MULTI-evolve FCNNs on DMS datasets with tens to hundreds of thousands of variants, when the eventual MULTI-evolve engineering tasks train on only hundreds of variants—a profoundly reduced sampling regime.

For completeness, in Appendix A we review standard results about the additive null model that we use as a baseline in the sequel. Briefly, each combinatorial variant is represented as a binary vector *x* ∈ {0, 1} ^*p*^ indicating which of *p* possible mutations are present, and the measured phenotype—typically log fold change with respect to wild-type [20–24]—is modelled as *y* = *β*^⊤^*x* + *ϵ* with additive Gaussian noise. Rather than naively using only single-mutant measurements as estimates of the additive effect vector *β*, we performed regression on all available training variants (singles, doubles, etc.) [22, 25–28], using the ridge estimator (5).

## Results

Our analysis uses the MULTI-evolve code [29]; data for APEX, dCasRx, and HuABC2 variant libraries [15]; and DMS benchmark datasets [30] from the supplementary materials of [1]. Additive models were fit using scikit-learn [31]. Code to reproduce our findings is available on GitHub.^1^

### MULTI-evolve’s multimutant engineering recapitulates an additive model

We consider the three engineering applications of MULTI-evolve presented in [1]: engineering the APEX peroxidase for improved activity, the dCasRx RNA-targeting CRISPR effector for improved RNA binding, and the HuABC2 anti-CD122 antibody for improved expression and binding. For APEX, single mutants were nominated by a PLM and experimentally characterized, and then doubles were formed from the top 15 single mutations and experimentally characterized. For dCasRx, single mutants were nominated by DMS, and then doubles were formed from the top 15 single mutations and experimentally characterized. For HuABC2, single mutants were again nominated by a PLM, and doubles were formed from the top mutations across the VH and VL chains; both expression and binding phenotypes were measured. These single+double mutant datasets were used to train MULTI-evolve’s FCNNs to ostensibly learn epistatic interactions. The trained FCNNs were then used to generate predictions for all possible 5–9-mutant APEX variants, all 5– 7-mutant dCasRx variants, and 3–7-mutant HuABC2 variants; highly ranked variants were selected for production and experimental characterization and found to be hyperactive. For HuABC2, variant selection was guided by the expression–binding Pareto frontier. This capability to engineer hyperactive multimutants is the central engineering result of the paper, and is attributed to the FCNN’s capacity to learn epistatic interactions and synergistic effects.

To evaluate this claim, we generated predictions for the same combinatorial spaces using the same training data, but using the ridge estimator (5) of an additive model instead of the FCNN (with log_2_ fold change with respect to the wild-type as the training target). For APEX, we trained on the 75-min Amplex-Red timepoint, matching the (−)A134P engineering campaign in [1]. For HuABC2, we fit separate additive models for expression and binding. We also regenerated predictions across the same combinatorial variant spaces using the MULTI-evolve FCNN (using hyperparameters selected by best Spearman correlation in cross validation as in the original study; see Fig. 3, excluding the “0 Layers” option). Figure 1 shows the joint scatter of additive model predictions vs. MULTI-evolve FCNN predictions across the combinatorial variant space for APEX, dCasRx, and both HuABC2 phenotypes, with the variants selected for production highlighted.

**Figure 1:**
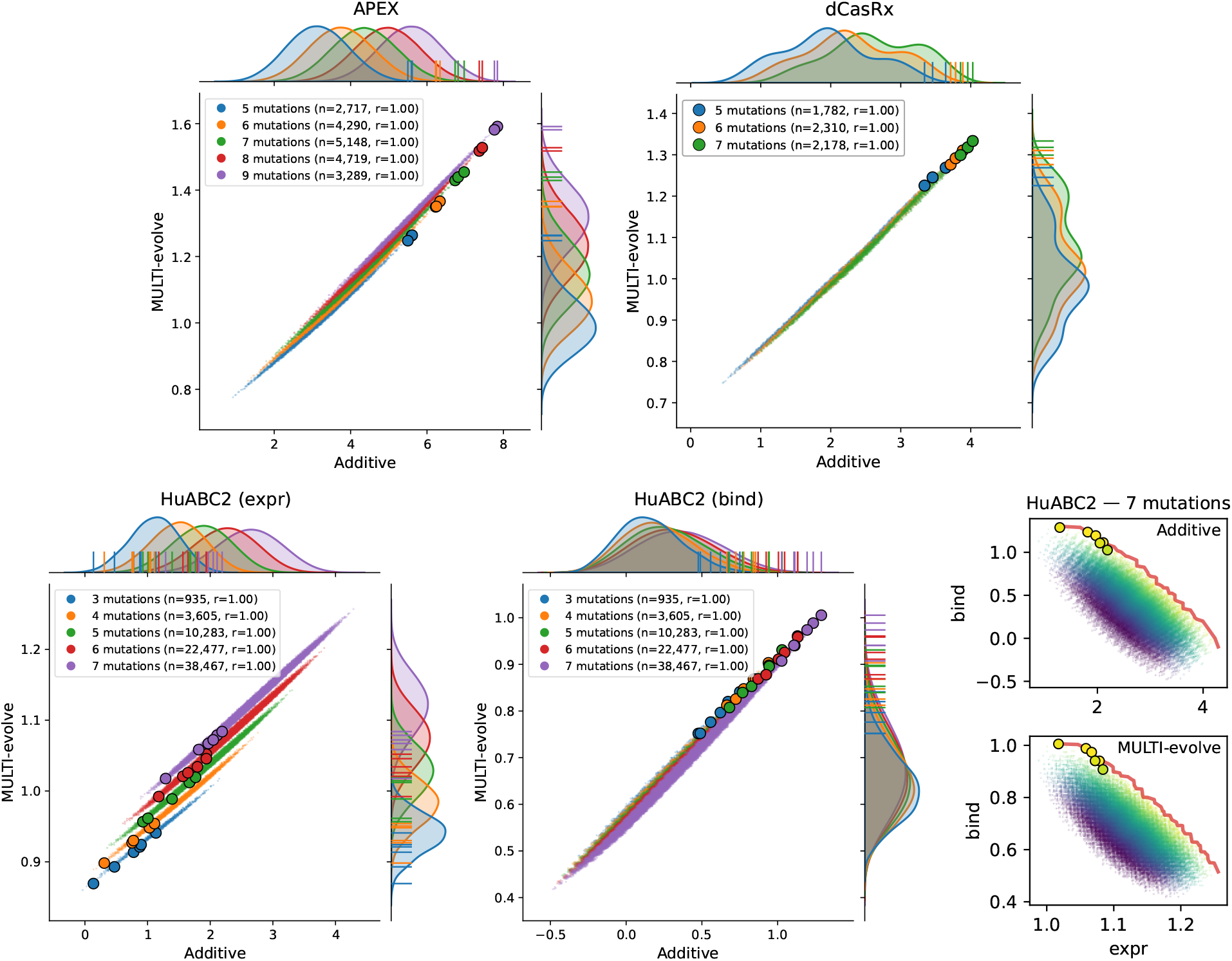
Additive model predictions vs. MULTI-evolve FCNN predictions across the combinatorial variant spaces. Joint scatter of additive vs. MULTI-evolve predictions, with per-group kernel density estimates in the marginal axes. MULTI-evolve predictions were made using as input to the NN the one-hot encoding of each protein’s amino-acid sequence. Each point is one combinatorial variant, colored by mutation count; larger outlined markers mark the variants selected for production in [1], with their locations also indicated as ticks on the marginal axes. The legend in each panel reports the variant count *n* and Pearson *r* for each mutation class. **Top row:** APEX (−)A134P (5-layer NN; 5–9 mutations) and dCasRx (1-layer NN; 5–7 mutations). **Bottom row:** HuABC2 expression (5-layer NN; 3–7 mutations), HuABC2 binding (3-layer NN; 3–7 mutations), and the expression–binding Pareto frontier for 7-mutation HuABC2 variants under the additive model (top) and MULTI-evolve (bottom). Small points show the full combinatorial cloud colored by distance from the additive model’s Pareto frontier; the red line traces the Pareto frontier; outlined markers indicate variants selected for production in [1]. The same additive-frontier distance is used to color points in both panels. See Fig. S1 for mutation counts 3–7 and Fig. S2 for alternative additive predictors.

We found that the predictions of the FCNN and the additive model are almost perfectly correlated across the combinatorial variant space for APEX, dCasRx, and both HuABC2 phenotypes (*r* > 0.999 in all cases), such that the ranking of tens of thousands of untested variants is practically identical to an additive model. In particular, the multimutant variants selected for production in [1] are similarly very highly ranked by both the FCNN and the additive model. For HuABC2, the expression–binding Pareto frontiers under the two models are nearly identical (Fig. 1, bottom right), meaning the multi-objective variant selection strategy is achieved using an additive model for each phenotype (not, as they claim, because MULTI-evolve “learned the distinct epistatic patterns governing each property and identified higher-order combinations that navigate around antagonistic interactions while amplifying beneficial ones”). These findings call into question the central claim that MULTI-evolve’s success is due to learning synergistic interactions and traversing a rugged epistatic landscape. The engineering strategy and outcomes are not distinguishable from the classical strategy of identifying beneficial single mutations and combining those with the largest additive effects. Additionally, in Appendix C we characterize the ordinary least squares (OLS) estimator (2) trained on singles+doubles with misspecification in terms of true additive and epistatic effects, showing that it is only powered to recover additive effects (true epistatic effects weakly influence estimated additive effects, and the influence grows weaker with larger singles+doubles training designs).

We also investigated whether MULTI-evolve represents any pairwise epistatic signal in its double-mutant training data. Figure 2 compares the FCNN-predicted epistatic residuals to the measured epistatic residuals for all doubles in the training set, for APEX, dCasRx, and both HuABC2 phenotypes. In all cases, the predicted residuals cluster tightly around a constant value of − *ŷ*_WT_, the value expected if the FCNN acts as a purely additive model with an intercept and learns no pairwise epistasis, and there is no correlation between the predicted and measured residuals. MULTI-evolve does not represent epistasis in the training data, and thus has no capacity to extrapolate it to higher-order mutants.

**Figure 2:**
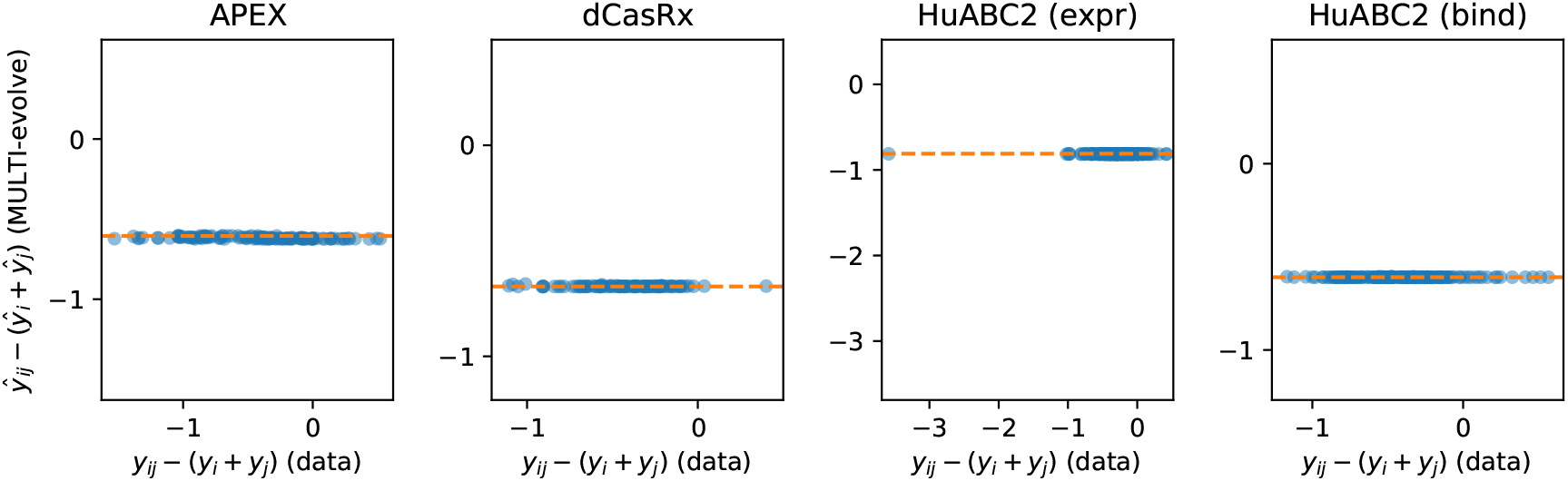
MULTI-evolve vs. measured epistatic residuals on training-set double mutants. Each point is one double mutant {*i, j*}. The measured epistatic residual *y*_*ij*_ − (*y*_*i*_ + *y*_*j*_) and the FCNN-predicted residual *ŷ*_*ij*_ − (*ŷ*_*i*_ + *ŷ*_*j*_), averaged over 10 cross-validation splits, are compared. The dashed line indicates − *ŷ*_WT_: if the FCNN recapitulates a linear model with an intercept term, all predicted residuals would fall on this WT intercept via the previous formula. From left to right: APEX, dCasRx, HuABC2 expression, HuABC2 binding.

### MULTI-evolve neural networks do not outperform additive models on held-out test data

The near-perfect agreement between the FCNN and the additive model on the combinatorial variant space raises the question of how the FCNN performs relative to an additive model on held-out test data from the original single+double training sets. The original study presents a hyperparameter grid search for the FCNN architecture, varying the number of hidden layers, the batch size, and the learning rate. The original study also varies the type of protein residue representation, but as the study consistently found that a simple one-hot encoding of the amino-acids types delivered the best validation performance, we only considered this representation in our analysis. We ran this grid search on APEX (−)A134P, dCasRx, and HuABC2 (expression and binding) data using the published code [29], modified to extend the grid to include zero hidden layers, which is tantamount to a linear model with no capacity to represent epistasis. We also added a classical ridge estimator (5) of an additive model as a baseline. Results are shown in Fig. 3. We find that, for APEX, dCasRx, and both HuABC2 phenotypes, the MULTI-evolve FCNNs do not significantly outperform the zero-hidden-layer linear model or the additive model (dashed lines) in terms of held-out test set performance.

**Figure 3:**
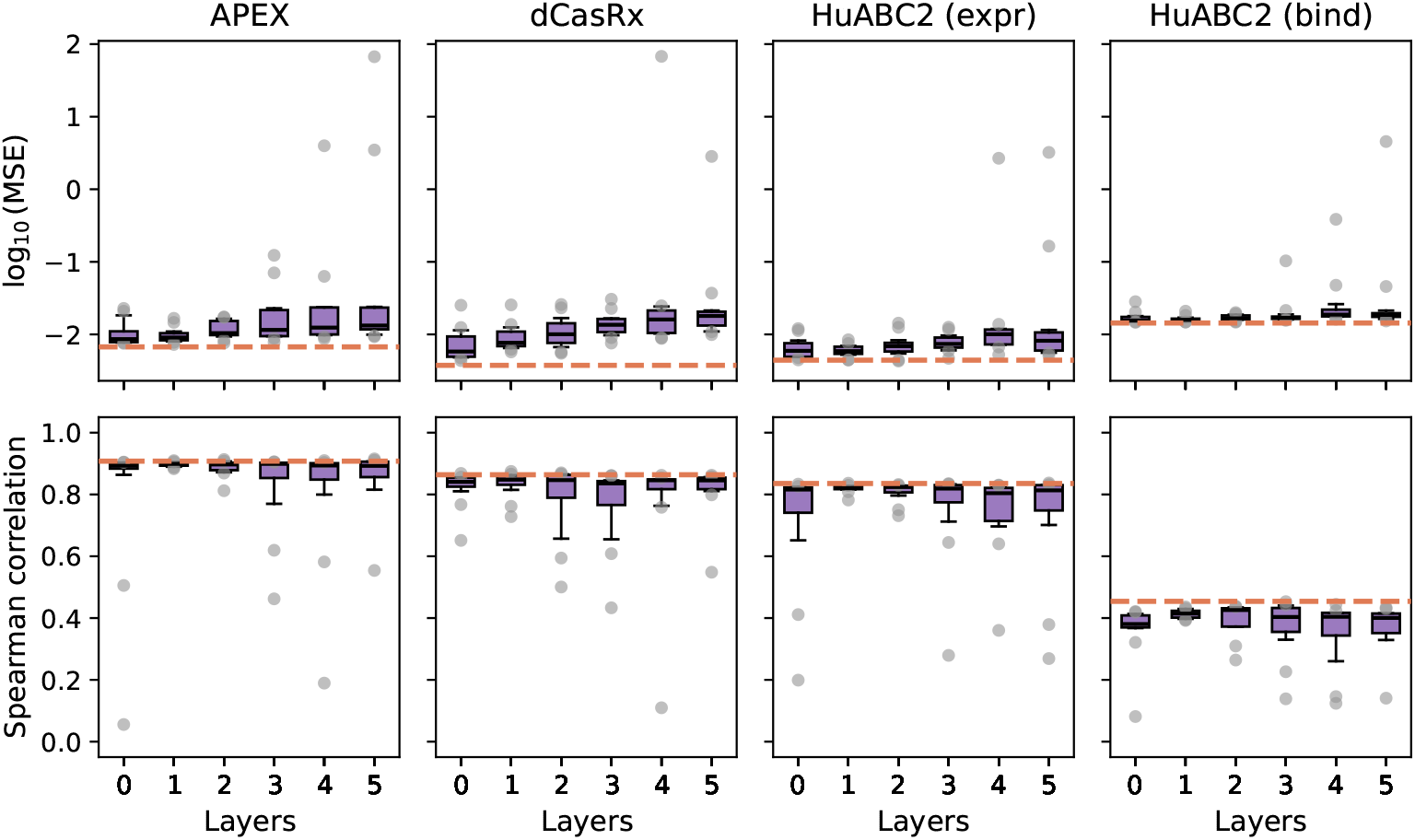
Held-out test performance across the MULTI-evolve hyperparameter grid. Boxplots show the distribution of test-set MSE **(top)** and Spearman correlation **(bottom)** across all configurations grouped by number of hidden layers, for APEX **(left)**, dCasRx **(second from left)**, and HuABC2 expression and binding **(right)**. We used MULTI-evolve’s hyperparameter sweep code, extending the grid to include zero hidden layers (tantamount to a linear model) and an additive model (ridge estimator (5), dashed orange line). Our grid also included batch size and learning rate as in the original study, but these are suppressed in the plot for simplicity.

### The DMS benchmark does not demonstrate epistasis learning

Having shown that the MULTI-evolve FCNN recapitulates an additive model in the combinatorial variant engineering space and doesn’t outperform an additive model on held-out test data, we now revisit the DMS benchmarking experiment presented as key evidence that MULTI-evolve’s FCNN learns epistasis. In [1], the FCNN was trained on 12 previously published deep mutational scanning datasets [14, 17–19], using three nested training sets: (a) single mutants only, (b) single and double mutants only, and (c) single, double, and triple mutants. The authors report that the Pearson correlation between predicted and measured phenotypes on a held-out multimutant test set improves with each training dataset expansion, and interpret this as evidence that the FCNN learns epistatic interactions from the double-mutant training data, which enables it to extrapolate to higher-order mutants (see their Fig. 2E). Statistical support is assessed by paired *t* tests comparing the distribution of Pearson *r* across the 12 datasets and 5 CV folds for each training set, showing significant improvement when doubles are added to singles, and further improvement when triples are added to singles and doubles. The authors write that “incorporating double mutants in the training set markedly improved predictive performance over single mutants alone” and interpret this as evidence that “double mutants capture sufficient epistatic information for accurate extrapolation to higher order mutants.”

The interpretation of this trend as evincing learned epistasis is unfounded. We show that the pattern of improved prediction when the training set is enlarged is expected even under an additive null model, due to the elementary statistical phenomenon of prediction error variance reduction with larger sample sizes. We further demonstrate that an additive model trained on the same nested training datasets reproduces the pattern in [1, Fig. 2E] on the same test set. Indeed, we find that the MULTI-evolve FCNN does not outperform this additive model in terms of test set prediction performance.

Under the additive model Eq. (1), suppose we fit on *p* single-mutant measurements only. Each mutation *i* appears in exactly one training observation, so OLS estimates are 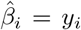 and have variance *σ*^2^ due to measurement noise. The predicted phenotype of a *k*-mutant is 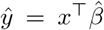, where *x* is the binary vector indicating which mutations are present. The prediction error variance for the singles-only design is Var [*ŷ* − *y*] = (*k* + 1) *σ*^2^, where the first term is the coefficient estimation error and the second is measurement noise in *y*.

Now suppose we add all 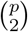 pairwise double mutants, and construct the OLS estimator on the combined design of 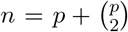 training observations. Under a null additive model, an elementary calculation from theory of linear regression (see Appendix B) yields the prediction error variance for the singles+doubles design, to first order in 1*/p*, as Var [*ŷ* − *y*] = (*k/p* + 1) *σ*^2^. The prediction error variance reduction of the singles+doubles design compared to the singles-only design with *p* mutations is given by the ratio for the two cases. For a large design *p* ≫ *k* this is approximately a 1/(*k* + 1)-fold reduction. Additionally, in Appendix C we generalize the null to consider an additive regression model applied to data generated from a true model with pairwise epistasis, and derive an analogous variance reduction result, as well as a misspecification bias term given as a design-weighted sum of the true pairwise epistatic effects.

Although the elementary statistical points above show that the [1, Fig. 2E] pattern is expected under an additive null model, one might object that this analysis doesn’t apply to the multimutant test set which likely contains pervasive epistatic effects that can’t be captured by the additive model, so the improvement in prediction when doubles and triples are added to training must reflect learning of those epistatic effects. To address this, we used the ridge estimator (5) to fit an additive model to the same nested training datasets used for the DMS benchmark, and evaluated the Pearson correlation between predicted and measured phenotypes on the same held-out multimutant test set. Figure 4 shows that the additive model reproduces the pattern of improved multimutant prediction when doubles and triples are added to training—the same pattern presented as evidence of epistasis learning in [1, Fig. 2E]. We also observed that the raw Pearson *r* values for the additive model are not systematically lower than those of MULTI-evolve, and in fact are higher for the singles-only training set (Fig. 4, bottom panels).

**Figure 4:**
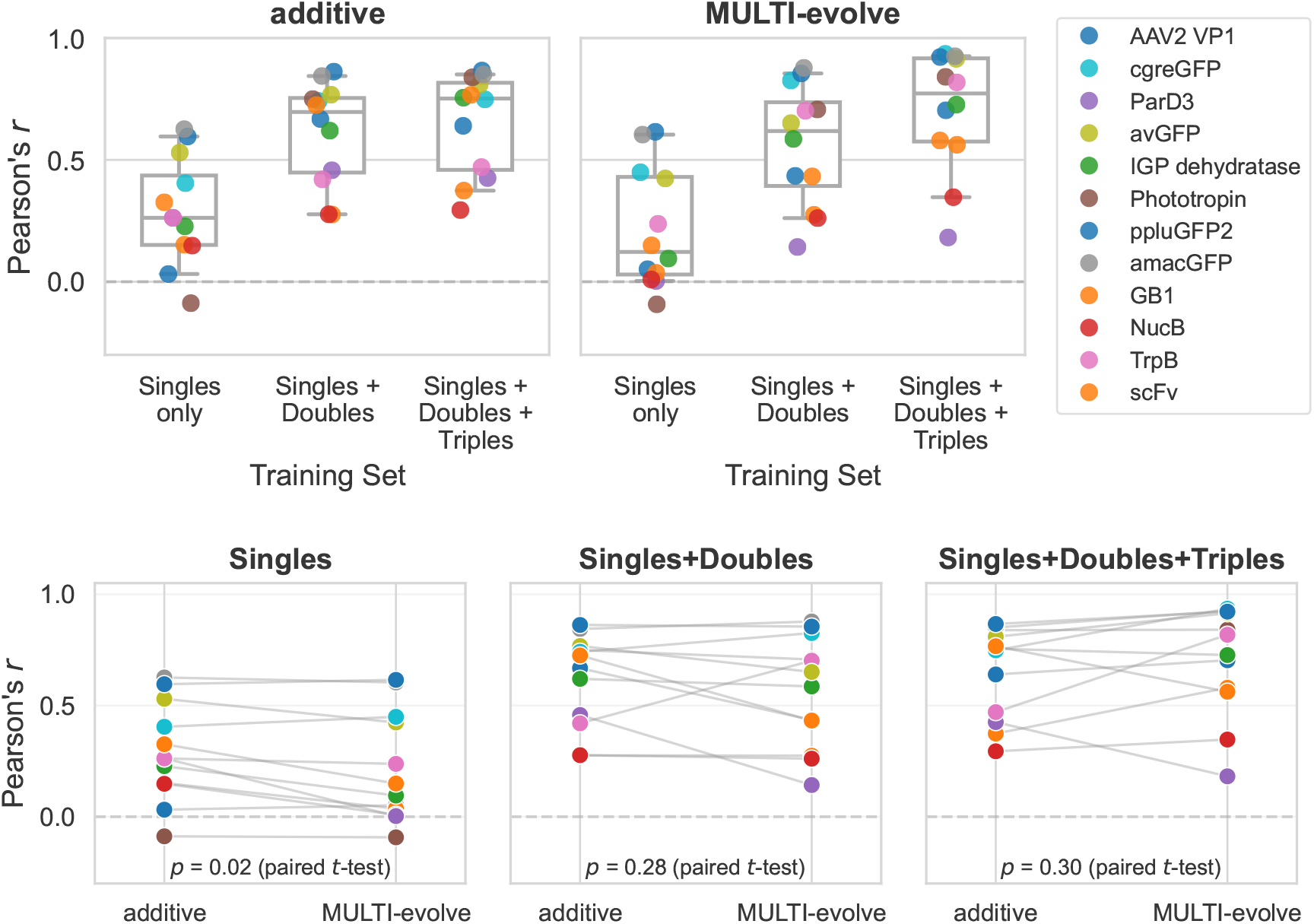
An additive model for the benchmark datasets reproduces reported signature of epistasis learning. We used the classical ridge estimator (5) to fit an additive model to the benchmark datasets [30]. **Top left panel** shows that training the additive model on successively larger nested training sets (singles only, singles + doubles, singles + doubles + triples) produces the same pattern of improved multimutant prediction as reported for MULTI-evolve in [1, Fig. 2E] (reproduced in the **right panel** using source data from [1, Table S3]). **Bottom panels** show paired comparisons of the additive model and MULTI-evolve on the same benchmarks.

Finally, we note that the benchmarking regime of the DMS datasets is profoundly different from the regime of the engineering applications of MULTI-evolve considered above. The DMS datasets contain tens to hundreds of thousands of variants, including thousands of doubles and triples sampled broadly across sequence space. By contrast, the MULTI-evolve engineering applications train on only ∼ 100–200 doubles formed from only the top ∼ 15 single mutations nominated by PLMs or functional screens—a tiny, highly biased corner of the combinatorial landscape. These are profoundly different sampling regimes, and whether a method benefits from doubles in the former regime says little about what it learns in the latter. Regardless, we find that the DMS benchmark results are consistent with an additive null model, and that MULTI-evolve does not outperform an additive model in this regime either.

## Discussion

Genotype-phenotype maps of proteins are complex and may contain pervasive epistatic interactions that confound classical engineering strategies based on additive mutation stacking. The promise of machine learning-guided directed evolution is that ML models can learn these interactions and thereby navigate rugged landscapes to engineer superior proteins. MULTI-evolve is presented as a framework that achieves this promise by learning epistatic interactions from limited double-mutant training data and extrapolating to synergistic higher-order combinations. Schematic figures and language throughout the paper (and associated media content) depict a rugged nonlinear landscape and suggest that MULTI-evolve is able to “jump” to hyperactive multimutants by identifying epistatic synergies. We have shown that this is misleading: MULTI-evolve’s predictions across the combinatorial APEX, dCasRx, and HuABC2 variant spaces are almost perfectly correlated with an additive model’s, such that the ranking of tens of thousands of untested variants is virtually identical to rankings from an additive model—including the multi-objective expression–binding Pareto frontier for HuABC2; the FCNN does not outperform an additive model on held-out test data from these engineering tasks; and the DMS benchmarking results presented in support of the epistasis learning claim are expected under a null additive model, which reproduces the reported pattern when fit to the same data. MULTI-evolve learns an additive model, not epistasis.

Richer baselines beyond the strictly additive model can be considered, in which apparent epistasis arises without pairwise interaction terms *per se*. For *global epistasis*, mutation effects combine additively on a latent molecular phenotype but are observed through a nonlinear link, such as a saturating experimental readout [22]. Another case of interest is where the measurement depends on interaction between two molecular phenotypes: for example, protein stability can buffer the functional effects of mutations, so that a mutation’s effect on catalytic activity or binding depends on a latent stability budget shared across mutations [32, 33]. In both cases, the underlying architecture may be additive at a molecular-phenotype level, with nonlinearity entering through the mapping to the measurement process. Such models may be useful as more nuanced baselines because they can capture apparent epistasis without positing *bona fide* pairwise (or higher-order) interactions among mutations, and thus can be used to contextualize and interpret claims of learning such interactions. In the present work, the strict additive baseline already matches MULTI-evolve’s performance, so there is no gap for these richer baselines to close.

There is considerable excitement in the field about the potential of ML models to learn complex genotype-phenotype maps and enhance protein engineering with ML-in-the-loop workflows. We share this excitement. However, in the interest of advancing knowledge, we also advocate for critical and rigorous approaches to evaluating claims in this space. Additive model baselines are particularly important for evaluating claims of epistasis learning, since they have no capacity to represent epistasis of any kind. More broadly across machine learning, it is standard practice to compare complex nonlinear models such as neural networks against simple linear baselines to establish that the former are learning nontrivial patterns [34, 35]. In a critique of methodological standards in the ML community, Lipton & Steinhardt [36] cataloged several patterns that undermine scientific claims in ML research, including the failure to perform ablations against simple baselines, the conflation of a model’s capacity to represent complex functions with evidence that it has learned them, and the misattribution of empirical gains to a paper’s headline contribution—typically a complex model—when the actual source of performance is more mundane. Each of these issues is evident in MULTI-evolve: the engineering success is real, but the source of performance is that mutational effects are sufficiently additive for the proteins and mutations considered, not that the neural network has learned epistatic synergies. The FCNN learns additivity but the authors misattribute its performance to epistasis because no linear baseline was tested, leading to claims of a major advance on a longstanding challenge— learning to navigate epistatic genotype-phenotype maps. In fact, to explain MULTI-evolve’s benchmarking performance and engineering results, additivity is all you need.

## Acknowledgements

WSD thanks Sarah Hilton, Tyler Starr, Jesse Bloom, Jakub Otwinowski, Bill Noble, Saori Sakaue, Yun Song, and Erick Matsen for discussions and feedback.

## Appendix A

**The additive null model**

We consider a protein with *p* potential single-amino-acid mutations relative to wild-type, and represent each combinatorial variant as a binary (one-hot) vector *x* ∈ {0, 1} ^*p*^ indicating which mutations are present. The additive model posits that the measured phenotype of variant *x* is additive over the effects of each mutation and contains additive Gaussian noise:

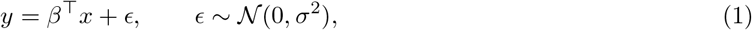

where *β* ∈ ℝ^*p*^ denotes the vector of additive effects of each mutation, and *σ*^2^ is the variance of the measurement noise. While a naive additive model approach would be to directly take single-mutant measurements as estimates of *β*, a more general and more standard approach is to fit the additive model using all available training variants (singles, doubles, triples, etc.) via regression [22, 25–28]. Ordinary least squares (OLS) regression solves

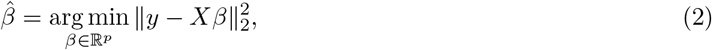

where 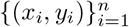 are the training observations and the *n × p* design matrix *X* has rows 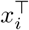. When *X* has full column rank, the OLS solution is given by the closed-form expression

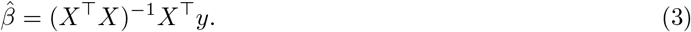

Ridge regression adds an *ℓ*_2_ penalty to the coefficients, which can be helpful when the design matrix is not full rank or is ill-conditioned, solving

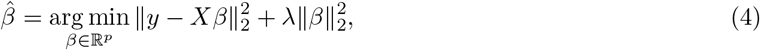

where *λ >* 0 is a regularization hyperparameter (usually determined via cross-validation). This has the closed-form solution

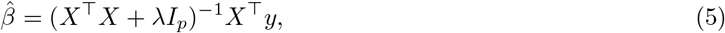

where *I*_*p*_ denotes the *p × p* identity matrix.

## Appendix B

**Prediction error variance reduction with singles+doubles design**

The variance-covariance matrix of OLS estimates is in general given as 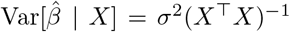, so we need to compute (*X* ^⊤^ *X*)^−1^ for the combined design (singles and doubles). Each double {*i, j*} contributes one training observation in which both *x*_*i*_ = 1 and *x*_*j*_ = 1. In the combined design, each mutation *i* appears in 1 single and *p* − 1 doubles, so the diagonal of *X* ^⊤^ *X* has entries 1 + (*p* − 1) = *p*. The off-diagonal entries are all 1 (each pair of mutations co-occurs in exactly one double). Applying the Sherman-Morrison formula [37], we find

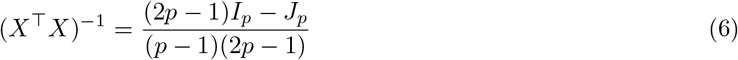

where *J*_*p*_ denotes the *p × p* matrix of all ones. The diagonal entries give the variance of each 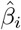 as

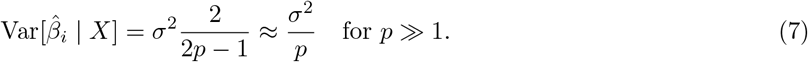

The off-diagonal entries give the covariance between estimated mutation effects 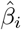 and 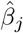 as

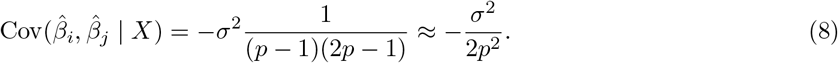

The predicted phenotype of a *k*-mutant is 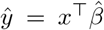, where *x* is the binary vector indicating which mutations are present. The prediction error variance for the singles-only design is

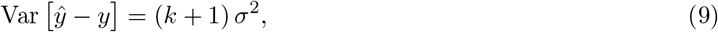

where the first term is the coefficient estimation error and the second is measurement noise in *y*. For the singles+doubles design, the prediction error variance is, to first order in 1*/p*

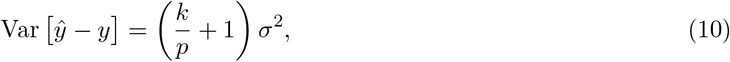

where we have used Eqs. (7) for the first term.

## Appendix C

**Influence of pairwise epistasis on additive model regression with the singles+doubles design**

We continue the OLS analysis of Appendix B, now obtaining a closed-form expression for 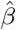 in terms of hypothetical true pairwise epistatic effects. Reusing (*X* ^⊤^ *X*)^−1^ from Appendix B and noting that 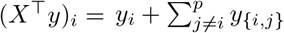, the OLS solution 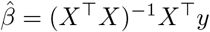 is

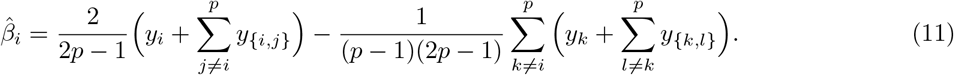

In the above, *y*_*i*_ denotes the measured phenotype of the single mutant with mutation *i*, and *y*_{*i,j*}_ denotes the measured phenotype of the double mutant with mutations *i* and *j* (note mutations are unordered, so that *y*_{*i,j*}_ = *y*_{*j,i*}_). This expression becomes interpretable once we decompose phenotypes into signal and noise. For single mutants, *y*_*i*_ = *β*_*i*_ + *ϵ*_*i*_, where *β*_*i*_ is the true mutation effect and *ϵ*_*i*_ ∼ *N* (0, *σ*^2^) is i.i.d. measurement noise. For doubles, we decompose into an additive part, a true pairwise epistatic effect *β*_{*i,j*}_, and noise: *y*_*{i,j}*_ = *β*_*i*_ + *β*_*j*_ + *β*_{*i,j*}_ + *ϵ*_{*i,j*}_ (this decomposition does not assume any specific form of pairwise epistasis). Substituting into Eq. (11):

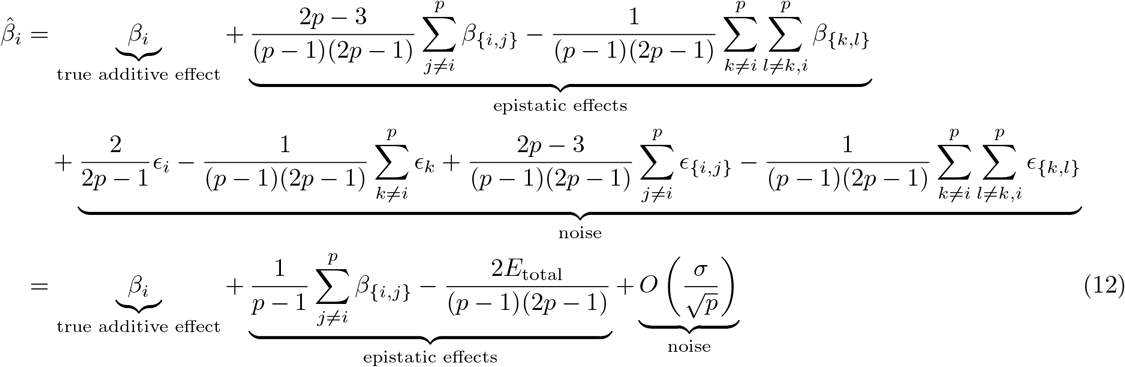

where 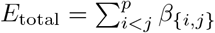 is the sum of all pairwise epistatic effects, and the noise term has variance *O*(*σ*^2^*/p*), recovering the result of Appendix B that adding doubles sharply reduces estimator variance relative to using single-mutant measurements directly (Var = *σ*^2^). This alone produces a boost in test-set prediction even in the complete absence of epistasis, contributing to the trend in Fig. 4. When true pairwise epistasis is present, the OLS estimates of additive effects are additively biased by averages of the true pairwise epistatic effects. Because the additive model is underspecified, this represents a bias trade-off between single- and double-mutant predictions whose effect on higher-order test-set prediction depends on the structure of the epistatic landscape, and may further contribute to the trend in Fig. 4. Indeed, the prediction that the additive model makes on a variant with *k* mutations is

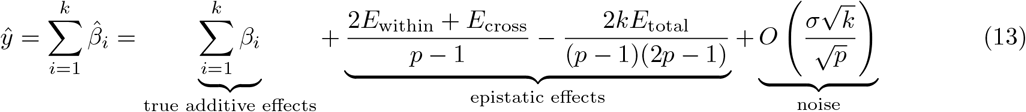

where *E*_within_ = ∑_1≤*i<j*≤*k*_ *β*_*{i,j}*_ contains all the epistatic effects between pairs of mutations in the *k*-mutant, and 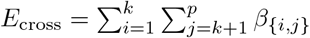 contains all the epistatic effects between one mutation in the *k*-mutant and one outside of it.

Again, depending on the structure of the epistatic landscape, the epistatic effects in the additive model’s predictions (Eq. 13) may contribute to the trend in Fig. 4. However, we note that these epistatic effects only approximate a small portion of the relevant epistatic signal, making the additive model’s predictions rely most heavily on its low-variance estimates of the true additive effects. Indeed, a true model of pairwise epistasis would be

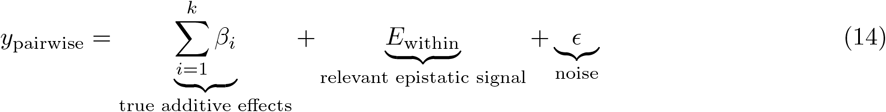

thus the additive model only recovers a biased estimate of 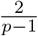 of the relevant epistatic signal, with bias coming from the *E*_cross_ and *E*_total_ terms.

## Supplementary Figures

**Fig. S1:**
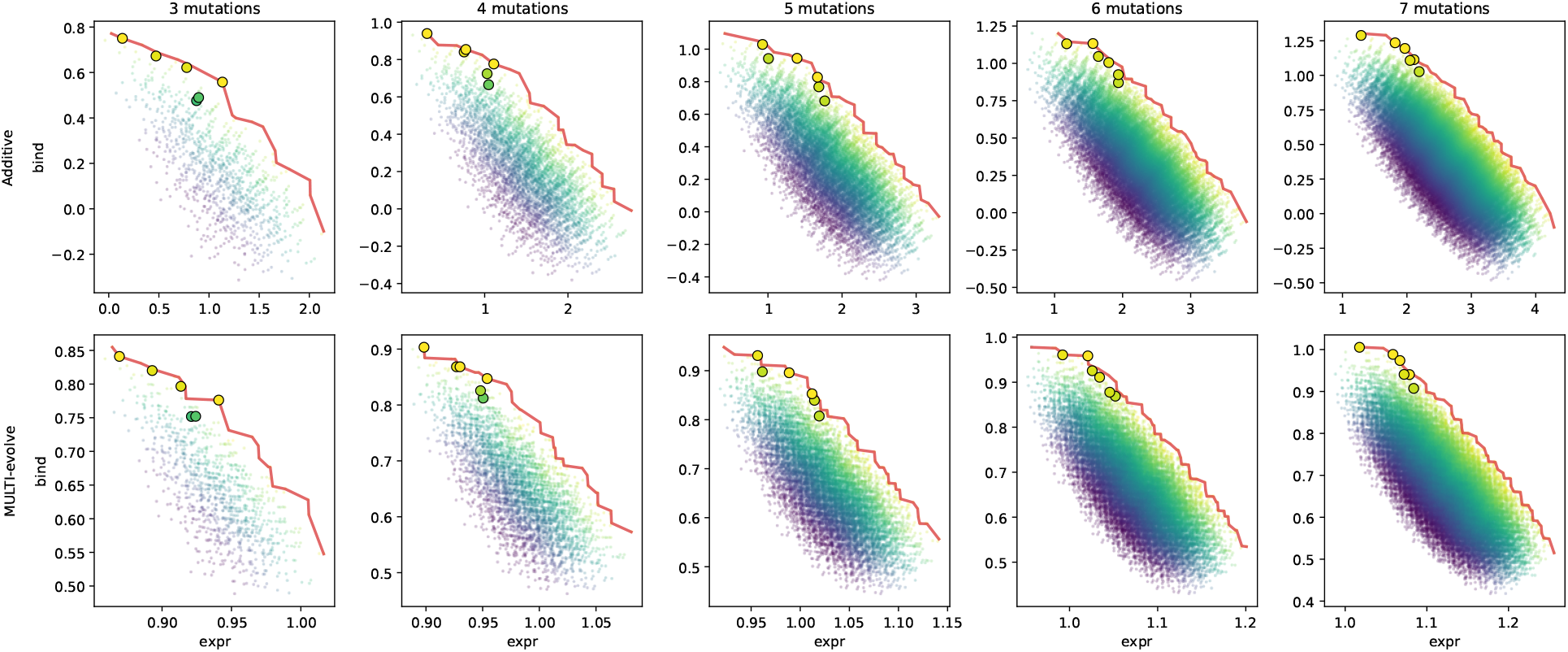
Expression–binding Pareto frontiers for HuABC2 across mutation counts 3–7. As in Fig. 1 (bottom right), but for all mutation counts 3–7.

**Fig. S2:**
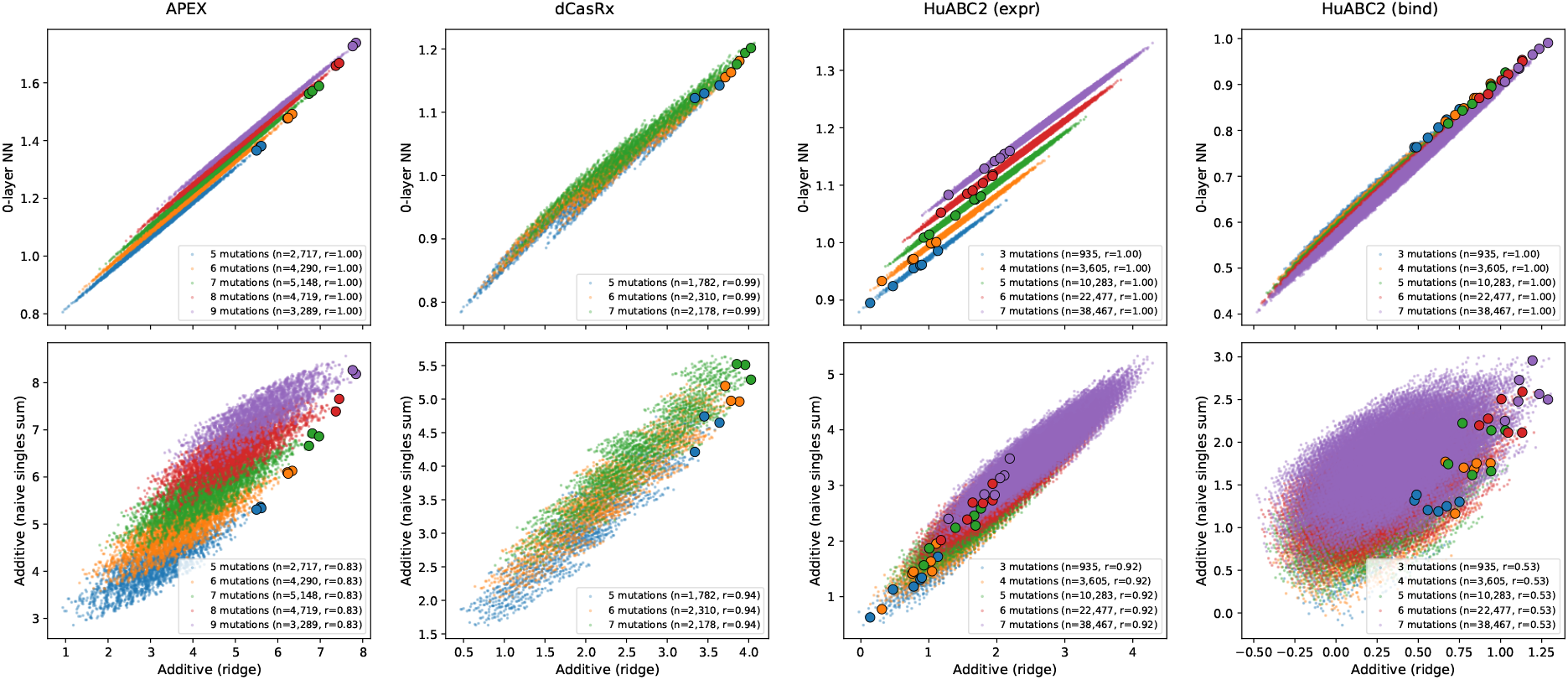
Alternative additive predictors agree with the ridge estimator on the combinatorial variant space. **Top row:** APEX (left) and dCasRx (right). **Bottom row:** HuABC2 expression (left) and HuABC2 binding (right). Within each protein panel: ridge additive predictions (*x*) vs. 0-layer MULTI-evolve predictions (*y*) (upper), and ridge additive predictions (*x*) vs. a naive additive predictor that adds measured single-mutant effects (*y*) with no regression (lower). Points are colored by mutation count; variants selected in [1] are outlined in black. The legend reports *n* and Pearson *r* per mutation class.

## Footnotes

1 https://github.com/dewitt-lab/multievolve-baselines

